# *Cis* element length variability does not confer differential transcription factor occupancy at the *D. melanogaster* histone locus

**DOI:** 10.1101/2024.06.24.600460

**Authors:** L.J. Hodkinson, L.E. Rieder

## Abstract

Histone genes require precise regulation to maintain histone homeostasis and ensure nucleosome stoichiometry. Animal histone genes often have unique clustered genomic organization. However, there is variability of histone gene number and organization as well as differential regulation of the histone genes across species. The *Drosophila melanogaster* histone locus has unique organizational characteristics as it exists as a series of ∼100 highly regular, tandemly repeated arrays of the 5 replication-dependent histone genes at a single locus. Yet *D. melanogaster* are viable with only 12 transgenic histone gene arrays. We hypothesized that the histone genes across the locus are differentially regulated. We discovered that the GA-repeat within the *H3/H4* promoter is the only variable sequence across the histone gene arrays. The *H3/H4* promoter GA-repeat is targeted by CLAMP to promote histone gene expression. We also show two additional GA-binding transcription factors, GAGA Factor and Pipsqueak, target the GA-repeat. When we further examined CLAMP and GAF targeting, we determined that neither CLAMP nor GAF show bias for any GA-repeat lengths. Furthermore, we found that the distribution of GA-repeats targeted by both CLAMP and GAF do not change throughout early development. Together our results suggest that the transcription factors targeting the *H3/H4* GA-repeat do not impact differential regulation of the histone genes, but indicate that future studies should interrogate additional *cis* elements or factors that impact histone gene regulation.

## Introduction

Histone genes need to be strictly regulated so there are neither too few nor too many histones at any given time in the cell. The canonical histone genes, *H3, H4, H2A, H2B,* and *H1*, are replication dependent; their regulation is strictly coupled to the cycle. Histone genes are expressed during S phase to package newly replicated DNA followed by halting of the gene expression by the end of G2. In part due to the requirement for strict coordinated cell cycle regulation, animal histone genes often have unique clustered genomic organization; however, there is variability of histone gene organization and differential regulation across species.

Histone genes were originally cloned and sequenced from purple and green sea urchin, *S. purpuratus* and *P. miliaris,* respectively, in the late 70s from which we learned sea urchin genomes have two sets of histone genes. The first set consists of a tandem repeat of the five canonical histone genes termed the “early histone genes’’ and a second set of 39 genes that are separated from the early genes, termed the “late histone genes” (Marzluff *et al*. 2006). These two histone gene sets are differentially regulated based on cell type and timing. The early histone genes are only expressed in the egg through the blastula stage whereas the late histone genes are expressed during late embryogenesis and continue expression through adulthood in all somatic cells (Marzluff *et al*. 2006).

The human genome also carries two clusters of histone genes, a major cluster on chromosome 6 and a minor cluster on chromosome 1. All H1 genes are located in the major cluster which is spread across several megabases and contains ∼60 histone genes in smaller sub-clusters while the minor cluster only contains around 10-12 histone genes (Seal *et al*. 2022; Ghule *et al*. 2023). Recent Hi-C data shows there are distinct promoter-promoter interactions between the subclusters of the major histone locus on chromosome 6, which suggest regulatory mechanisms could be different between the major and minor locus (Carty *et al*. 2017; Ghule *et al*. 2023). Transcription factors that regulate histone genes are shared between these loci, however the loci are differentially regulated to ensure there are correct stoichiometries of H3, H4, H2A, H2B and H1. The major histone locus also associates with the Cajal body throughout the cell cycle whereas the minor locus only associates with it during S phase (Ma *et al*. 2000; Shopland *et al*. 2001). From work in human embryonic stem cells, the H4 genes may have distinct regulation patterns between he major and minor loci based on tumor cell type. However, the patterns of H4 gene expression show only minor differences between loci in embryonic stem cells. This suggests that the overall contribution of histone transcripts from the major and minor loci are similar, implying that there are mechanisms of differential regulation that maintain this equilibrium despite differences in histone gene copy number between the loci (Becker *et al*. 2007).

Fission yeast are an even more extreme example of how histone genes are differentially regulated. Fission yeast genomes contain three pairs of H3-H4 genes, along with just a single pair of H2Aalpha-H2B, and a lone H2Abeta gene. A study investigating the three pairs of H3-H4 genes found that the first and third pair are up-regulated while the second pair is normally downregulated, exhibiting oscillation of expression through the cell cycle (Takayama and Takahashi 2007).

The histone genes in *Drosophila melanogaster* are a unique example of clustered histone gene organization. The *D. melanogaster* genome carries a single repetitive histone locus on chromosome 2L. Based on recent locus assembly from long-read sequencing (Bongartz and Schloissnig 2019), the *D. melanogaster* histone locus includes approximately 100 tandemly repeated histone gene arrays. Each 5kb array includes the five canonical histone genes, *H1, H3, H4, H2A* and *H2B,* along with their respective *cis* regulatory elements and promoters. *H3* and *H4* share a bi-directional promoter that contains an important GA-repeat *cis* element critical for histone gene expression (Salzler *et al*. 2013; Rieder *et al*. 2017; Hodkinson *et al*. 2024). *D. melanogaster* can survive with a 12-array histone transgene (Günesdogan *et al*. 2010; Salzler *et al*. 2013; McKay *et al*. 2015) indicating that not every gene is necessary. The studies from yeast, sea urchin, and even humans led us to hypothesize that individual histone genes or groups of genes are differentially regulated based on developmental timing, gene copy number, and number of loci.

To explore how the arrays at the endogenous *D. melanogaster* histone locus might be functionally different, we utilized a recent histone locus assembly completed through long-read sequencing (Bongartz and Schloissnig 2019) to search for sequence differences between the histone gene arrays. We discovered that the arrays are nearly identical in sequence, but the GA-repeat in the s *H3/H4* promoter is variable in length ranging from 16-35 nucleotides. The *H3/H4* promoter sequence can nucleate recruitment of specific histone regulatory factors, and the GA-repeat is specifically targeted by the transcription factor CLAMP (Salzler *et al*. 2013; Rieder *et al*. 2017; Koreski *et al*. 2020). Further, we recently confirmed that the GA-repeats are critical for histone locus factor recruitment (Rieder *et al*. 2017; Hodkinson *et al*. 2024). Therefore, we hypothesized that histone genes might experience differential regulation thorough variability of the GA-repeat and transcription factor occupancy.

To test this hypothesis, we obtained existing ChIP-seq data from CLAMP and other GA-repeat binding factors, GAGA Factor (GAF) and Pipsqueak (Psq), and investigated their differential occupancy over the histone GA-repeats. We discovered that all three factors bind the range of GA-repeat lengths and, furthermore, show that CLAMP and GAF are unbiased in the GA-repeat lengths they bind. Our discovery of variable GA-repeats at the histone locus uncovered a previously unknown distinction of the ∼100 histone gene arrays and may provide a target for future studies on histone array uniqueness and functionality. Furthermore, our observations suggest that the GA-repeat variability likely does not contribute to differential occupancy of transcription factors at histone gene arrays and implies other *cis* elements or cofactors that might contribute to differential histone gene regulation.

## Results

### The GA-repeat is variable in length across the histone gene arrays

Bongartz *et al*. (2019) produced a *de novo* assembly of the *Drosophila melanogaster* repetitive histone gene locus, identifying that the locus contains ∼107 histone gene arrays, in alignment with previous estimates (Lifton *et al*. 1978; McKay *et al*. 2015) and recently confirmed by another group (Shukla *et al*. 2024). We aligned the gene arrays (**Figure 1A**) and discovered that they are nearly identical in sequence other than length variability of a GA-repeat present in the bidirectional promoter of genes *H3* and *H4* (**Figure B**). The GA-repeat varies in length from 16 base pairs to 35 base pairs (**Figure 1B, C**). The most common GA-repeat length is 21 bp (29 of the 107 arrays). We found some clustering of arrays with similar length GA-repeats such as those that have GA-repeats with 29 or 31 bp GA-repeats (**Figure 1D**).

**Figure 1:**
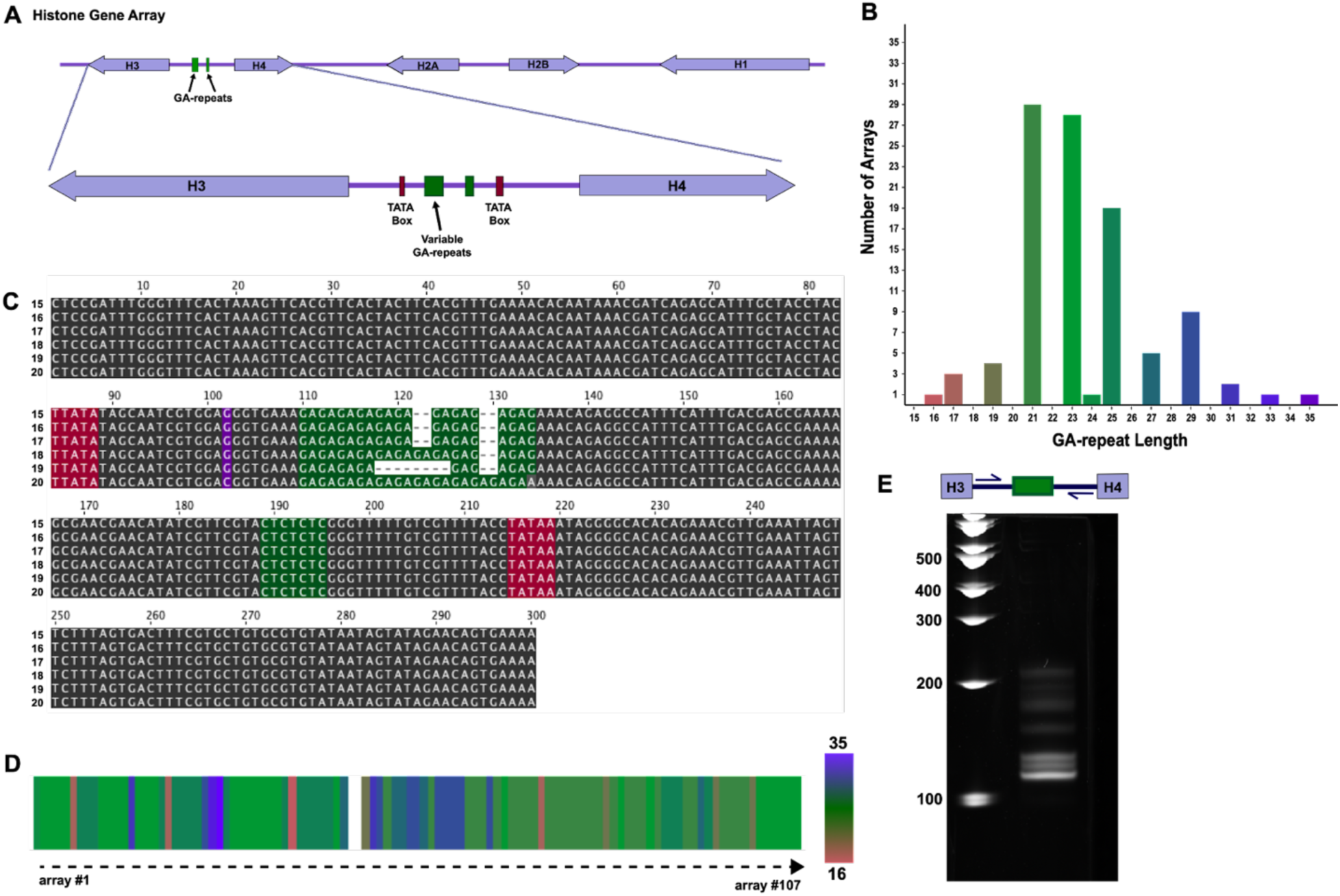
The GA-repeat length is variable across the histone gene arrays. **(A)** A diagram of a single histone gene array and the GA-repeat located in the *H3/H4*p. **(B)** We utilized previously assembled histone locus from Bongartz *et al*. (2019) to compare the sequences of the histone gene arrays. The arrays are virtually identical other than the GA-repeat in the *H3/H4*p, which varies in length. **(C)** We aligned six of the 300 bp *H3/H4*p (arrays 15-20, TATA boxes in maroon). Other than a single SNP (purple), the GA-repeat remains the only sequence variability. **(D)** A heatmap shows the positions of different GA-repeat lengths across the locus. Each array is represented by one vertical bar. **(E)** We designed primers to amplify the *H3/H4*p of the histone arrays to confirm the variability of the GA-repeat *in vivo*. Laddering of PCR products in an acrylamide gel confirmed GA-repeat variability.

To confirm the GA-repeat variability *in vivo*, we designed primers to amplify 115 bp of the endogenous *H3/H4*p region that includes the GA-repeat region. PCR from genomic DNA is predicted to produce amplicons ranging from 110 bp (16 bp GA-repeat) to 129 bp (35 bp GA-repeat). We observed the expected laddering of PCR products on an acrylamide gel, confirming GA-repeat length variability *in vivo* (**Figure 1E**). We noticed several amplicons that exceeded the predicted length, possibly due to secondary structure forming due to the GA-repeat.

### CLAMP, GAF, and Psq all target the GA-repeats in the *H3/H4*p

Our observations indicate that the most dramatic sequence difference across the histone arrays is the wide variability of the GA-repeat length. We previously demonstrated that this sequence is targeted by the CLAMP transcription factor (Rieder *et al*. 2017) and that the interaction is important for HLB factor recruitment and histone gene expression (Rieder *et al*. 2017; Hodkinson *et al*. 2024). However, the *Drosophila* genome encodes two other GA-repeat binding transcription factors: GAGA Factor (GAF) and Pipsqueak (Psq) (Lehmann *et al*. 1998; van Steensel *et al*. 2003) (**Figure 2A**). We therefore hypothesized that these other GA-repeat binding factors also target the GA-repeats in the histone gene array. To test our hypothesis, we aligned previously generated ChIP-sequencing data to the histone gene array (Gutierrez-Perez *et al*. 2019; Gaskill *et al*. 2021; Duan *et al*. 2021).

**Figure 2:**
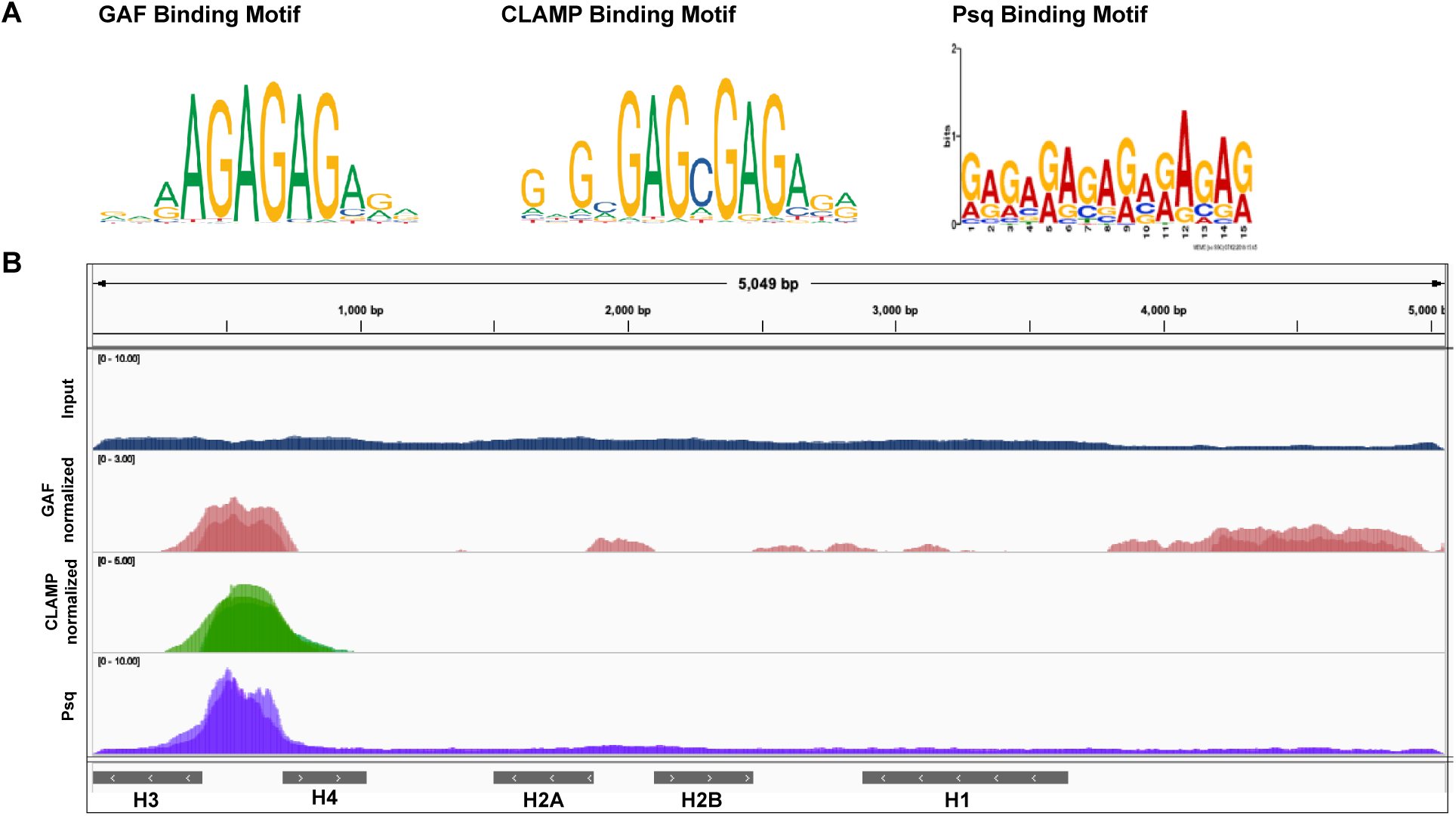
GAF, CLAMP, and Psq target the *H3/H4*p GA-repeat. **(A)** The binding motifs for GAF, CLAMP and Psq all contain GA-repeats. Binding motifs for GAF and CLAMP generated by the open access database JASPAR (Castro-Mondragon *et al*. 2022) and Psq binding motif recreated from Gutierrez-Perez *et al*. 2019. **(B)** We aligned ChIP-seq data for GAF (pink, two replicates overlayed (Gaskill *et al*. 2021)) in 2-3 hr embryos, CLAMP (green, three replicates overlayed (Duan *et al*. 2021)) in 2-4 hr embryos, and Psq (purple, two replicates overlayed (Gutierrez-Perez *et al*. 2019)) in 2-4 hr embryos to the single histone gene array. We normalized GAF and CLAMP data to respective inputs. We did not normalize Psq data because no inputs were provided in the original dataset. All three factors target the GA-repeat in the *H3/H4*p of the histone gene array. A representative input (blue) from the GAF and CLAMP ChIP-seq data is shown for comparison.

Because of the repetitive nature of the histone locus, aligning sequencing data such as ChIP-seq becomes impractical as each read would map to more than one or even all of the ∼ 100 arrays. Historically, to align sequencing data to the histone gene array, we utilized a condensed or custom version of the histone gene array, similar to the single histone gene array in McKay *et al*. (2015). Using the condensed histone gene array also means there is only one GA-repeat *cis* length (21 bp).

First, we confirmed that CLAMP robustly targets the *H3/H4*p GA-repeat **(Figure 2B).** CLAMP shows a clear peak over this region, as previously observed (Rieder *et al*. 2017; Koreski *et al*. 2020). Previously published GAF ChIP data from 2-3 hr embryos (Gaskill *et al*. 2021) and Psq data from *Drosophila* embryonic stem cells (Kc167 cells (Gutierrez-Perez *et al*. 2019)) also show a clear peak at the *H3/H4*p. Based on these data, CLAMP, GAF and Psq all target the histone locus. The factors may compete with each other or act synergistically.

### GA-binding factors do not show preference for GA-repeat length at the histone locus

Although the ChIP peaks shown in **Figure 2** for all three GA-repeat binding factors indicate that they target the histone genes, we cannot deduce which array(s) they target because we are only looking at the data aligned to a single histone array rather than the entire locus. We next wanted to deduce what arrays each of the three GA-repeat binding factors might target by examining whether they have a bias for certain length GA-repeats. Because the GA-repeats are the only sequence that differs between the ∼ 100 histone gene arrays, determining what length GA-repeats CLAMP, GAF, and Psq target can help us infer which arrays they target. CLAMP shows preference for binding longer GA-repeats on the X chromosome while GAF shows preference for short GA-repeats (Kaye *et al*. 2018). *In vitro*, CLAMP binds DNA probes with long GA-repeats up to 30 nucleotides in length by EMSA, whereas GAF will only shift probes with shorter GA-repeats of 8 nucleotides (Kaye *et al*. 2018). Therefore, we hypothesized that these GA-repeat binding factors might target different histone arrays based on their binding preference for GA-repeat length.

We developed a bioinformatics script that selected *H3/H4*p sequences from the ChIP-seq dataset by defining two anchor sequences, one upstream (5’) and one downstream (3’) of the GA-repeat with enough length to ensure specificity to the *H3/H4*p. The code then extracts the reads that match both anchors, scans to identify the GA-repeat and counts the number of nucleotides that make up the GA-repeat in that read. We utilized the ChIP input as a positive control, hypothesizing that we would confirm the GA-repeat lengths and frequencies we retrieved from the long-read sequencing results (**Figure 1B**).

When we generated histograms for GA-repeat length frequency from the input libraries of 0-2 hr and 2-4 hr embryos, we observed that the distribution of GA-repeat lengths mirrored the distribution we found from the long read Bongartz *et al*. (2019) data (**Figure 1B**). However, we did notice a few differences. We identified some GA-repeats that were shorter than expected due to SNPs in the middle of the GA-repeat (**Supplementary Figure 1**). In addition, we noticed some minor differences in the frequencies of element lengths (**Figure 1B** vs. **Figure 3**). This is likely due to genotype, as the Bongartz assembly was obtained from OregonR *Drosophila*, while the ChIP-seq datasets were produced from *yellow-, white-* maternal triple GAL-4 driver (MTD-GAL4, Bloomington, #31777) *Drosophila* (Ni *et al*. 2011). Because large, repetitive regions of the genome are subjected to frequent expansion and contraction due to unequal crossing over (Smith 1976; Shukla *et al*., 2024), few *Drosophila* strains have exactly the same GA-repeat length distribution (Shukla *et al*., 2024). Even individuals within an interbreeding population may have different numbers of arrays and therefore frequencies of GA-repeat lengths. Overall, however, we confirmed that the variability and length distribution of the histone locus GA-repeats is relatively reproducible.

**Figure 3:**
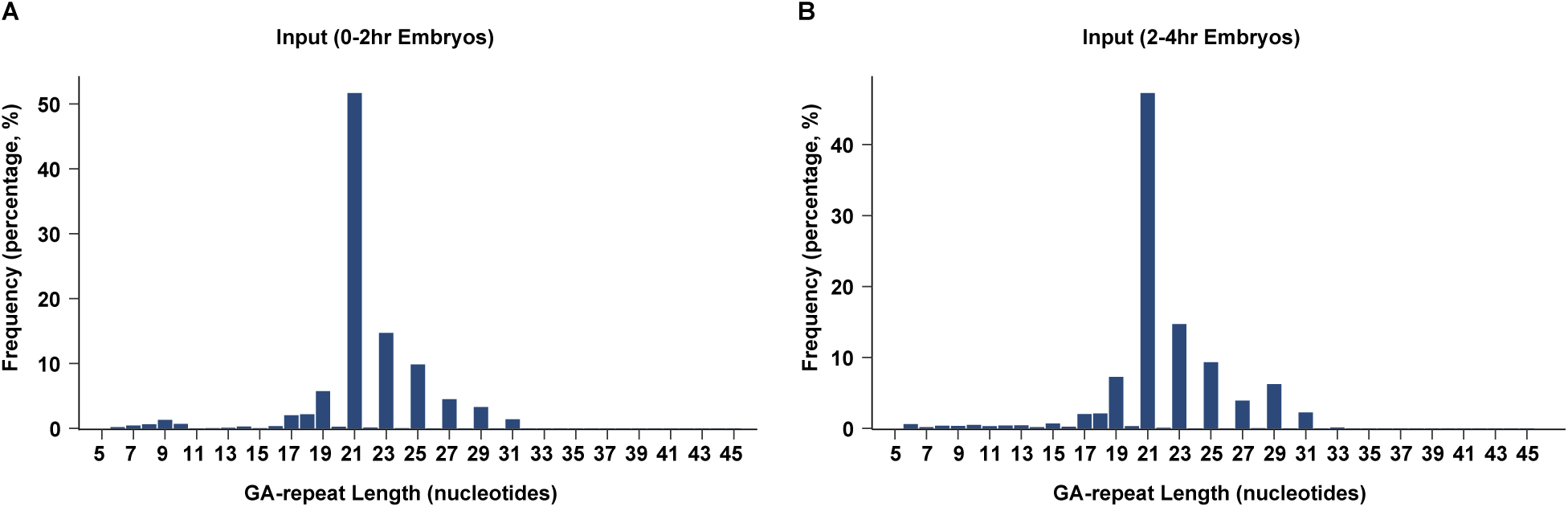
The *H3/H4* promoter GA-repeat length variability is observed in different datasets. We designed a bioinformatics code to parse ChIP-seq datasets, extract reads containing the *H3/H4* promoter GA-repeat and count the number of nucleotides within the repeat. We extracted reads from input ChIP-seq datasets of **(A)** 0-2 hr embryos (three replicates) and **(B)** 2-4 hr embryos (three replicates) and created histograms based on the GA-repeat lengths. The X-axis shows all GA-repeat lengths, and the Y-axis is the frequency each length was found represented as a percentage of extracted reads which contained that GA-repeat length. Data from Duan *et al*. 2021.

To determine the binding profiles of CLAMP and GAF at the variable GA-repeat, we used an available CLAMP ChIP-seq dataset from Duan *et al*. (2021) and generated a GAF ChIP dataset, both of which include data from 0-2 and 2-4 hr *Drosophila* embryos. These are relevant time points for histone gene expression; the early *Drosophila embryo* undergoes 14 nuclear division cycles during which the entire genome is replicated every 8-12 minutes. Therefore, a large number of histones are rapidly required (Tadros and Lipshitz 2009; Farrell and O’Farrell 2014; Harrison and Eisen 2015). The histone genes are targeted by specific factors as early as nuclear cycle 9 (Terzo *et al*. 2015), and zygotic histone genes are expressed by nuclear cycle 11 (Edgar and Schubiger 1986). CLAMP is maternally deposited and targets the histone locus in the early embryo, prior to detectable histone gene expression (Rieder *et al*. 2017). GAF is not thought to target the zygotic histone locus unless CLAMP is depleted (Rieder *et al*. 2017), although we discovered that it likely does so, at least from some datasets (**Figure 2**).

Using these embryonic ChIP-seq datasets, we investigated the frequencies of GA-repeat lengths targeted by CLAMP and GAF. We found that all GA-repeat lengths were bound by CLAMP, which does not show any bias for specific GA-repeat lengths despite preferring long X-linked GA-repeats (Kaye *et al*. 2018). Furthermore, CLAMP seems to target each of the GA-repeat lengths at similar frequencies to their respective counts across the locus (**input, Figure 3**). Lastly, we found no difference between the distribution of GA-repeat lengths targeted by CLAMP based on age of embryo, suggesting that developmental timing does not impact CLAMP occupancy at the histone locus GA-repeats (**Figure 4C, D**).

**Figure 4:**
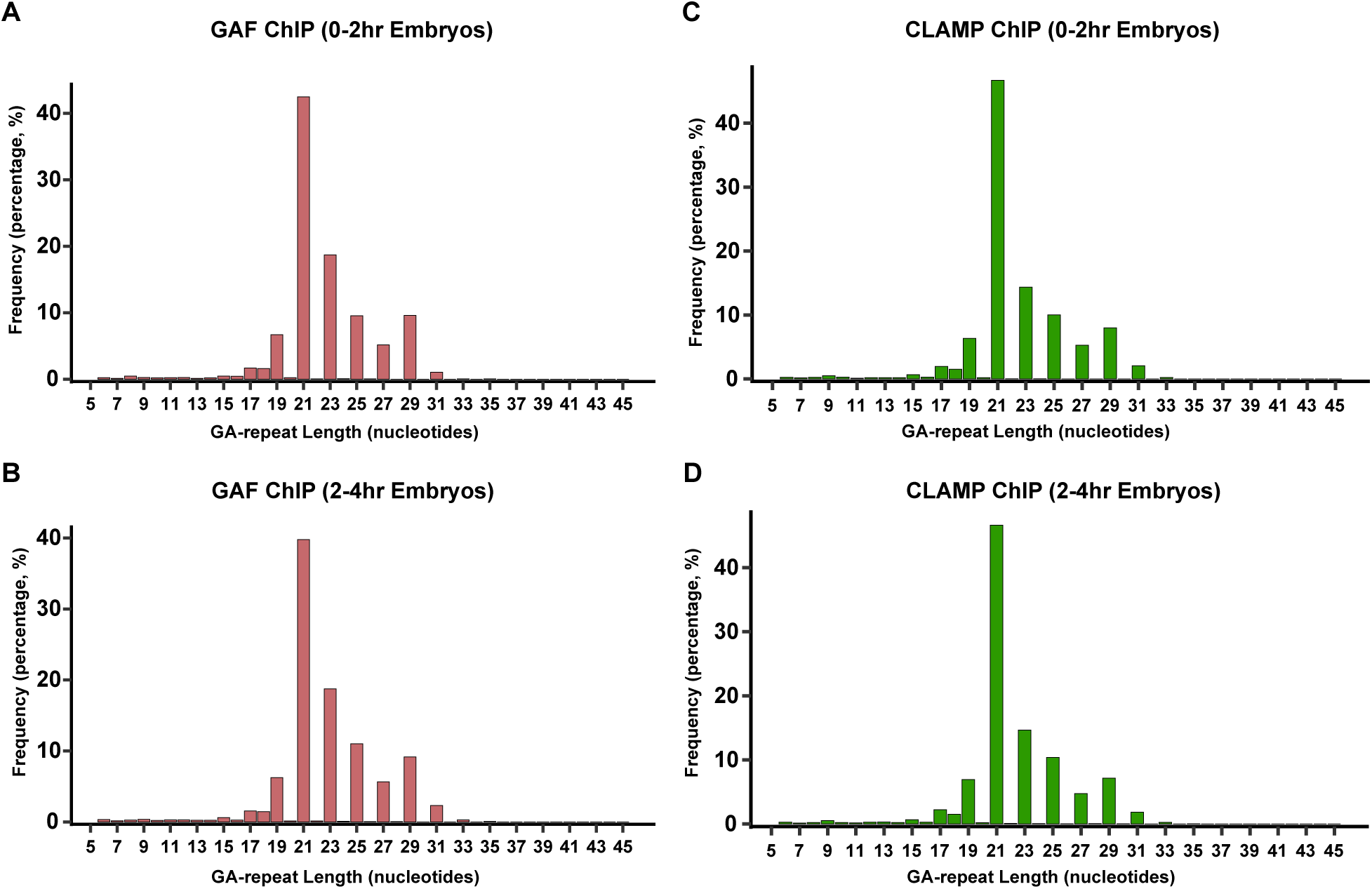
GAF and CLAMP target the same length GA-repeats. We extracted reads containing the *H3/H4* promoter GA-repeat from ChIP-seq data for GAF in **(A)** 0-2 hr embryos (three replicates) and **(B)** 2-4 hr embryos (three replicates). We also extracted reads contain the *H3/H4* promoter GA-repeat from ChIP-seq data for CLAMP in **(C)** 0-2 hr embryos (three replicates) and **(D)** 2-4 hr embryos (three replicates). In all histograms, the X-axis shows GA-repeat lengths, and the Y-axis is the average frequency each length was found represented as the percentage of extracted reads.

We next performed a similar analysis for GAF and retrieved similar results. GAF showed no bias for specific GA-repeat lengths in either 0-2 or 2-4 hr embryos (**Figure 4A, B)**. Further, we found that GAF also seems to bind each of the GA-repeat lengths at similar frequencies to their respective counts across the locus (**input, Figure 3**) similar to CLAMP (**Figure 4C, D**). Although we identified a previously generated Psq ChIP-seq dataset from 3 hr embryos, we were unable to interrogate GA-repeat binding preference due to the short length (50 bp) of the sequencing reads, which is not sufficient to identify reads that contain both the anchor sequences and the GA-repeat.

## Discussion

The *Drosophila melanogaster* histone locus comprises ∼100 virtually identical histone gene arrays and is regulated by a unique nuclear body. It is unknown whether all ∼100 histone genes are all targeted by the same transcription factors and produce the same mRNA output, as we are unable to map histone transcripts to their genes of origin. Some evidence points toward differential expression of genes. Animals carrying 12-array transgenes in the background of an endogenous locus deletion are viable (Günesdogan *et al*. 2010; McKay *et al*. 2015; Zhang *et al*. 2019) and express histone mRNAs at the same level as the endogenous locus, indicating that 100 genes are not required for viability. Other species, including other drosophilids, have varying numbers of histone genes in differing genomic arrangements, such as the closely related *D. simulans* whose genome carries only 15 histone arrays (unpublished) or ∼40 MYa diverged *D. virilis* whose genome carries two histone loci which combined only contain 32 arrays (Russo *et al*. 1995; Schienman *et al*. 1998; Xie *et al*. 2022). It is difficult to assay how the *D. melanogaster* genes might be differentially expressed, as the histone coding sequences are virtually identical. We therefore sought to uncover mechanisms for differential histone gene regulation by investigating sequence differences between arrays.

Using a recent long-read histone locus assembly (Bongartz *et al*. 2019), we discovered that the GA-repeat in the *H3/H4* promoter is variable in length across the histone locus, while the rest of the 5 kb arrays are nearly identical in sequence. We previously demonstrated that CLAMP targets the GA-repeat in the *H3/H4* promoter and confirmed that the GA-repeats are important for histone locus factor recruitment (Rieder *et al*. 2017; Hodkinson *et al*. 2024). These observations indicated the importance of the GA-repeat in overall histone gene regulation, so we hypothesized that this sequence variability might be functionally important in recruiting different transcription factors. We found that all GA-binding transcription factors target the element, but that none seems to have a bias for longer or shorter repeats.

GA dinucleotide repeats are fairly common in many genomes and serve a variety of functions. Short tandem repeats (STRs) are commonly found in core promoter sequences to serve as targets for transcription factors or pioneer factors like GAF and CLAMP (Duan *et al*. 2021), which displace nucleosomes to ready the gene for transcription. Recent work looking at a conserved GA-repeats in the core promoter of early human embryonic development genes shows that differences in GA-repeat length at these genes can cause differences in expression levels (Valipour *et al*. 2013). These data confirm that GA-repeat length itself is sufficient to drive differential gene expression and, furthermore, may imply that the length of the GA-repeat in the histone gene arrays may also impact differential expression of the histone gene, even if this is not due to changes in CLAMP, GAF, or Psq binding.

GA dinucleotide repeats can also serve as insulators. GAF binding at GA-repeats is critical for insulation between genes and unrelated, neighboring enhancer sequences (Lehmann 2004; Gaskill *et al*. 2021). In mice, GA-repeat motifs within the Hox gene clusters are nucleosome-free and, when GAF targets these regions, chromatin boundaries are established to create domains so the Hox genes are insulated from their neighboring regulatory elements (Srivastava *et al*. 2013). Similarly, in *D. melanogaster* GAF localizes to the *Fab-7* boundary element from the Hox genes *Ubx, Abd-A* and *Abd-B*. GAF can target GA-repeats at the *Fab-7* element, which can determine its function as an insulator at different developmental time points (Schweinsberg *et al*. 2004). It is possible that the GA-repeat in the histone array acts as an insulator and, although it is located within the *H3/H4* promoter, it may serve multiple functions as the target for binding factors at a subset of arrays and as an insulator for others to modulate the expression if histone genes in different arrays.

Studies in sea urchins, which have two clusters of histone genes that are differentially regulated, show that the specific downregulation of the “early” H2A genes is regulated by an upstream (5’) GA-repeat serving as an insulator (Di Caro *et al*. 2004). This study suggests that the regulation of individual histone genes can be governed by *cis* elements. Furthermore, this data emphasizes that dinucleotide repeats, and specifically GA-repeats, have important functions across species and in many genomic contexts. Our observations suggest the GA-repeat may not impact differential expression of the *Drosophila* histone genes, however here we did not explore if GA-repeat length impacted individual histone gene expression levels. The repetitiveness of the histone locus makes it impossible to assess the expression of individual histone genes because there is little coding sequence. Future experiments could leverage a “barcoded” 12-array transgene where silent mutations in the histone coding sequences allow determination of differential expression from each gene. Using this system, we could investigate how GA-repeat length impacts histone gene expression rather than just GA-repeat binding factor occupancy.

GA-repeats are only one *cis* element that can affect gene expression and it is likely that there are secondary or several additional *cis* elements that are responsible for regulating histone genes (Horton *et al*. 2022; Hodkinson *et al*. 2023). The work here focused specifically on the GA-repeat as it is important for CLAMP binding and the only variable sequence between the arrays. However, additional *cis* elements in the *H3/H4* or *H2A/H2B* promoter could impact differential gene regulation. Furthermore, it is possible that all the arrays are targeted by GA-repeat binding factors, but additional transcription factors then activate the genes.

Here, we only consider the occupancy of the GA-repeat binding factors in the histone arrays, yet a body of factors regulates histone gene expression, known as the histone locus body (HLB) (Duronio and Marzluff 2017), the full composition of which is still unknown. Future studies exploring what other DNA-binding factors target the histone arrays, like the recently published screen from Hodkinson *et al*. 2024, as well as investigating how differential targeting may impact histone gene expression, will provide greater understanding of the intricacies involved in histone gene regulation.

By utilizing the previously assembled histone locus sequencing data (Bongartz and Schloissnig 2019), we revealed the *H3/H4* promoter GA-repeat *cis* element is the only variable sequence between the ∼100 gene arrays. We leveraged previously published ChIP-sequencing datasets and determined that the variability in the GA-repeat does not impact the occupancy of GAF and CLAMP and suggests that these factors alone are not responsible for any differential regulation of the histone genes. Overall, our results have expanded our understanding of the sequence features of the *D. melanogaster* histone locus and given insight into histone gene regulation mechanisms.

## Methods

### Promoter Alignment

We obtained the *H3/H4* promoter sequences from the Bongartz *et al*. (2019) genome assembly (**Figure 2A**) and used reads extracted from input ChIP-sequncing data (**Supplemental Figure 2**). We aligned sequences using T-Coffee Multiple Sequence Alignment (Notredame *et al*. 2000) to create a ClustalW output and formatted the shading and features with Jalview (Waterhouse *et al*. 2009).

### ChIP-analysis and Data Visualization - IGV plots

We directly imported individual FASTQ datasets into the web-based platform Galaxy (The Galaxy Community 2022) through the NCBI SRA Run Selector by selecting the desired runs and utilizing the computing Galaxy download feature. We retrieved the FASTQ files from SRA using the “Faster Download and Extract Reads in FASTQ format from NCBI SRA” Galaxy command. Because the ∼100 histone gene arrays are extremely similar in sequence, we do not utilize the dm6 or dm3 genomes and instead collapse ChIP-seq data onto a single histone array. We used a custom “genome” that includes a single *Drosophila melanogaster* histone array similar to that in McKay *et al*. (2015), which we directly uploaded to Galaxy using the “upload data” feature, and normalized using the Galaxy command “NormalizeFasta” specifying an 80 bp line length for the output .fasta file. We aligned ChIP reads to the normalized histone gene array using Bowtie2 (Langmead and Salzberg 2012) to create .bam files using the user built-in index and “very sensitive end-to-end” parameter settings. We converted the .bam files to .bigwig files using the “bamCoverage” Galaxy command in which we set the bin size to 1 bp and set the effective genome size to user specified: 5000 bp (approximate size of l histone array). If an input dataset was available, we normalized ChIP datasets to input using the “bamCompare” Galaxy command in which we set the bin size to 1 bp. We visualized the .bigwig files using the Integrative Genome Viewer (Robinson *et al*. 2011).

**Table 1.** ChIP-sequencing datasets. Specifics for the NCBI GEO datasets used including the GEO Accession number, the SRA Run selector numbers, the developmental time of each sample, and the cited source.

| Factor | GEO Accession # | SRA Run Selector | Developmental Timepoint | Citation |
| --- | --- | --- | --- | --- |
| **GAF**<br>GAGA<br>Factor<br>(Trl) | GSE152773 | **Anti GAF-GFP**<br>1 -SRR12045586<br>2 - SRR12045588<br>**Input**<br>1- SRR12045585<br>2 - SRR12045587 | 2-3 hr, stage 5 embryos | (Gaskill *et al.* 2021) |
| **CLAMP**<br>Chromatin<br>linked<br>adaptor | GSE152613 | CLAMP antibody<br>1 - SRR12024931<br>2 - SRR12024949<br>3 - SRR12024967<br>Input | 2-4 hr embryos | (Duan *et al.* 2021) |
| for MSL proteins |  | 1 - SRR12024933<br>2 - SRR12024951<br>3 - SRR12024969 |  |  |
| **Psq**<br>Pipsqueak | GSE118047 | PsqM (PsqTot) antibody<br>1- SRR7638403<br>2 - SRR7638404 | Kc167<br>*Drosophila* embryonic cell line | (Gutierrez-Perez *et al.* 2019) |

### ChIP-analysis and Bioinformatics Pipeline - GA-Repeat Histograms

Our annotated pipeline is available on GitHub (https://github.com/rieder-lab/count_GA) in the script entitled count_ag_repeats.py. We utilized packages SeqIO from biopython (Cock *et al*. 2009) to parse through fastq files and regex from anaconda or pip for all functional outputs. We also utilized logging from anaconda or pip to create a built-in log for the run as an informational output. We designed the code to first identify, and extract reads that contain the *H3/H4* promoter sequence by using two short, flanking “anchor” sequences to the left (5’) and the right (3’) of the GA-repeat (left sequence: TAGCAATCGT right sequence: CATTTCATTTGACGAGC). We used a counting mechanism to ensure that reads with both the left and the right anchor were extracted however there is also an information output for single matches. We then designed the code to scan through the extracted reads until encountering the specified string “AGAGAG” as a seed sequence for the GA-repeat. Once the GA-repeat is identified, we designed the code to count the number of nucleotides within the repeat. Of note, we designed the code to allow for 0 mismatches in the repeat which meaning repeats where two “A” nucleotides or two “G” nucleotides are adjacent to each other will only be counted until that “AA” or “GG” appears. We identified that there are a handful of GA-repeats that contains SNPs causing “AA” or “GG” stretches (**Supplementary Figure 1**). However, this feature of the pipeline is changeable to allow for any specified number of mismatches. The script outputs 6 files to a specified path destination. These outputs include a .tsv file with four columns of information; the first column is nucleotide count of the GA-repeat, the second column has the extracted repeat itself, and the third column with the trimmed read where the repeat originated, and the last column has the sequence ID (**Supplementary Figure 2**). This file allows confirmation of the GA-repeat nucleotide counts as well as access to the reads the pipeline extracted. The other 5 files are .fastq.gz files that include reads from the script parsing through the entire sequencing file which include: dual_match.fastq.gz containing all the full length reads that had both ancho sequences, left_only.fastq.gz containing reads that only matched the left anchor sequence, no_match.fastq.gz, containing reads that did not have either anchor sequence, right only.fastq.gz containing reads that only matched the right anchor sequence, and strange_match.fastq.gz contain reads with unexpected configurations of the anchor sequences such as forward and reverse complements of these sequences.

### ChIP Analysis – GA-Repeat Histograms

CLAMP ChIP-seq datasets from Duan *et al*. 2021 were retrieved from is deposited at NCBI GEO. The accession number is (GSE152598). GAF ChIP data was performed as described in Duan *et al*. 2021 with 10uL of GAF antibody (Fuda *et al*. 2015).

**Table 2.** ChIP-seq data used to generate GA-repeat length histograms.

| Target TF | Developmental Timepoint | GEO Accession # | SRA Run Selector # |
| --- | --- | --- | --- |
| Input | 0-2 hr embryo | GSE152613 | Input<br>1- SRR12024924<br>2 - SRR12024942<br>3 - SRR12024960 |
| Input | 2-4 hr embryo | GSE152613 | Input<br>1 - SRR12024933<br>2 - SRR12024951<br>3 - SRR12024969 |
| GAF | 0-2 hr embryo | pending | GAF antibody<br>pending |
| GAF | 2-4 hr embryo | pending | GAF antibody<br>pending |
| CLAMP | 0-2 hr embryo | GSE152613 | CLAMP antibody<br>1- SRR12024922<br>2 - SRR12024940<br>3 - SRR12024958 |
| CLAMP | 2-4 hr embryo | GSE152613 | CLAMP antibody<br>1 - SRR12024931<br>2 - SRR12024949<br>3 - SRR12024967 |

## Acknowledgments

We would like to thank the Rieder Lab for supporting this work. We would specifically like to thank Dr. Casey Schmidt for her invaluable insight into early experimental design and execution. We would also like to thank Dr. David Gorkin who wrote the Python pipeline and was an invaluable source of support and bioinformatics knowledge throughout this project. This work was supported by T32GM00008490 and F31HD105452 to LJH; and R00HD092625 and R35GM142724 to LER.

## Supplemental Figures

**Supplementary Figure 1:**
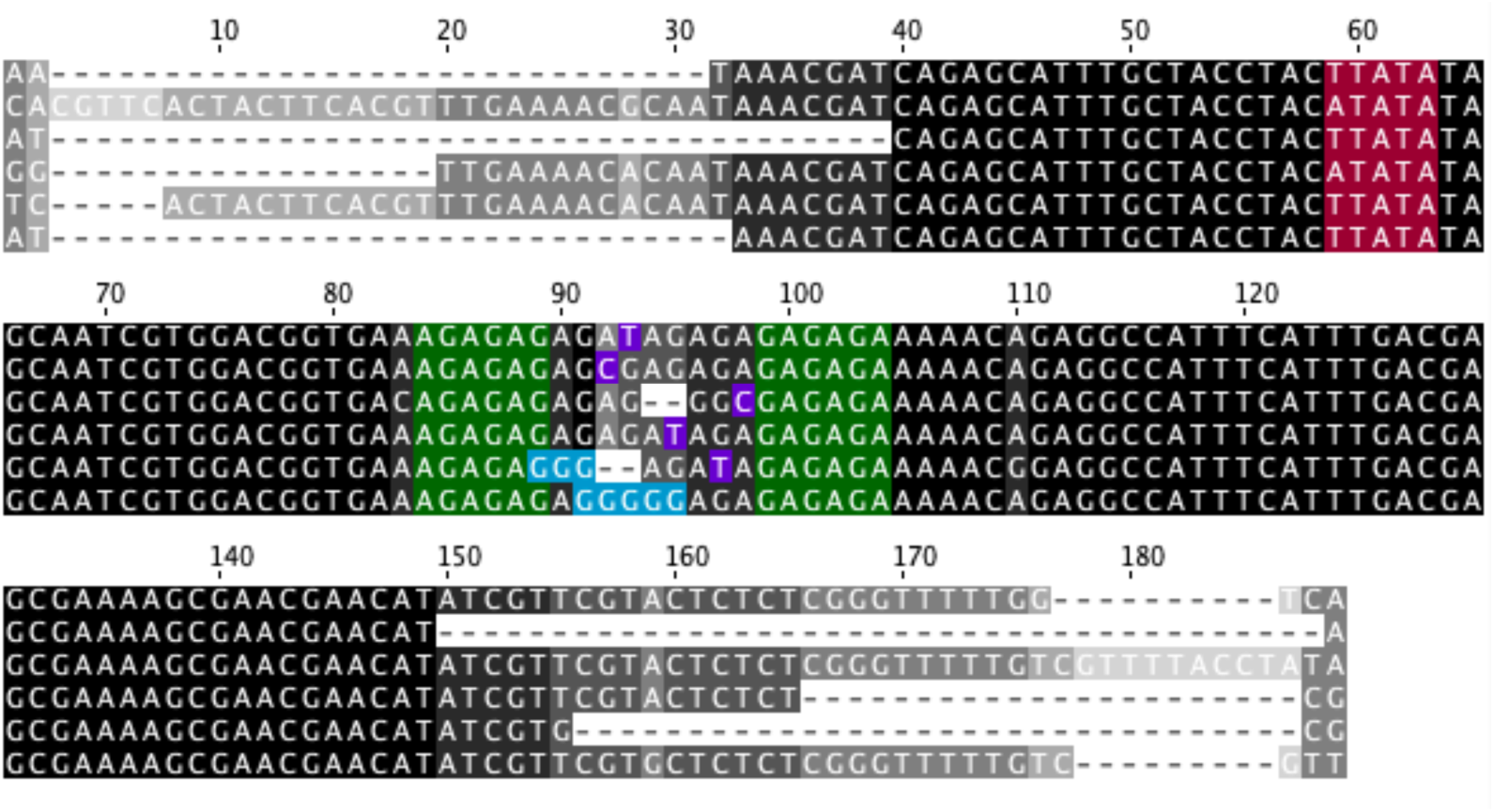
The GA-repeat can contain a variety of mismatches and sequence variation. We aligned a representative set of reads extracted by our bioinformatics pipeline where the *H3/H4* promoter GA-repeats contains SNPs or stretches of repeating A or G nucleotides. One of the TATA boxes is labeled in maroon and the GA-repeat is labeled in green. SNPs are shown in purple and stretches of A or G nucleotides are shown in teal. (Note these sequences have been extracted and trimmed by our Python script).

**Supplementary Figure 2:**
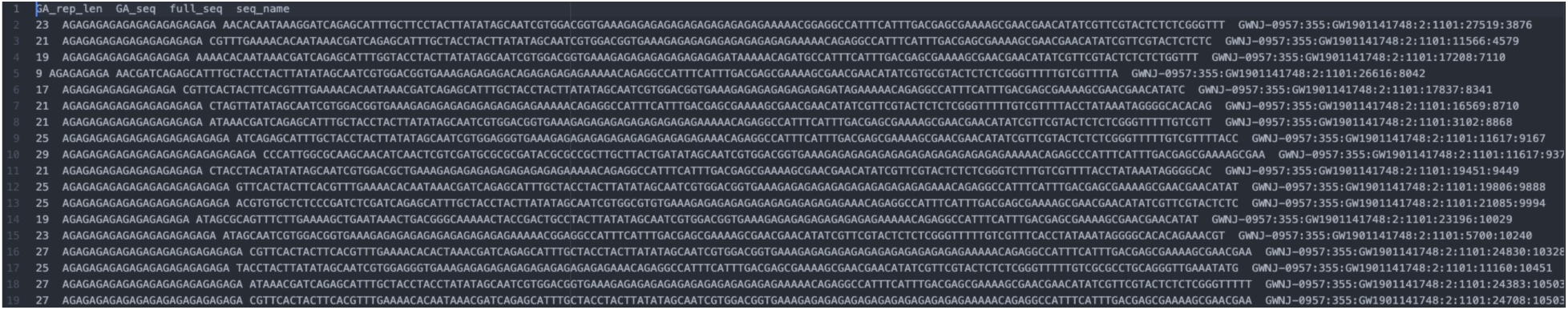
Sample .tsv file output for the GA-repeat counting bioinformatics pipeline. Our pipeline parses through sequence.fastq.gz files and extracts reads with the *H3/H4* promoter GA-repeat and counts the number of nucleotides that make up the repeat. The main output file for this script is a .tsv file containing four columns. The first column specifies the number of nucleotides that make up the GA-repeat, the second column has the extracted repeat itself, and the third column with the trimmed read where the repeat originated, and the last column has the sequence ID.

